# A Scalable, Multiplexed Assay for Decoding Receptor-Ligand Interactions

**DOI:** 10.1101/358739

**Authors:** Eric M. Jones, Rishi Jajoo, Daniel Cancilla, Nathan B. Lubock, Jeff Wang, Megan Satyadi, Rocky Cheung, Claire de March, Hiroaki Matsunami, Sriram Kosuri

## Abstract

Chemicals such as drugs, hormones, and odorants can have many potential interactions with endogenous targets, and uncovering these relationships is critical for understanding and modulating function. Mammalian olfactory receptors (ORs), a large family of G protein-coupled receptors, mediate olfaction through activation by small molecules. Each OR can respond to many odorants, and vice versa, making exploring this space one interaction at a time difficult. We developed a high-throughput receptor screening platform in human cell lines to screen libraries of chemicals against a multiplexed library of receptors using next-generation sequencing of barcoded genetic reporters. We screened three concentrations of 181 odorants, where in each well we record the activity of 39 ORs simultaneously, and identified 79 novel associations, including ligands for 15 orphan receptors. This platform allows the cost-effective mapping of large chemical libraries to receptor repertoires at scale.

---

Interactions between small molecules and receptors underpin an organism’s ability to sense and respond to its internal state and the environment. For many drugs and natural products, the ability to modulate many biological targets at once is crucial for their efficacy^1–3^. Thus, to understand the effect of many small molecules, we need to comprehensively characterize their functional interactions with biological targets. This many-on-many problem is laborious to study one interaction at a time, and is especially salient in the mammalian sense of smell^4,5^.

Olfaction is mediated by a class of G protein-coupled receptors (GPCRs) known as olfactory receptors (ORs)^6^. GPCRs are a central player in small molecule signaling and are currently targeted by 40% of US Food and Drug Administration (FDA) approved drugs^7^. ORs are a large family of class A GPCRs with approximately 396, 1130, and 1948 intact receptors in humans, mice, and elephants respectively^8^. Each OR can potentially interact with many odorants, and inversely, each odorant with many ORs. The majority of ORs remain orphan because of this vast combinatorial space, further compounded by the fact that recapitulating mammalian GPCR function *in vitro* is challenging^9,10^. In addition, no experimentally determined structure for any OR is available, hindering computational efforts to predict which odorants can activate each OR^11^.

Most GPCR and OR assays test chemicals against each receptor individually^12,13^. Multiplexed assays, where the activities of multiple receptors— often referred to as a library of receptors— are measured in the same well, would increase the throughput but have remained technically challenging. In such an assay, each cell expresses a single type of receptor and, upon activation, transcribes a short barcode sequence that identifies the particular receptor expressed in that cell. The enrichment of barcoded transcripts corresponding to each receptor’s activation are then measured by microarrays or next-generation sequencing. Such multiplexed GPCR activity assays have previously been attempted by transient transfection of individual receptors and subsequent pooled screening^14,15^. However, these assays are difficult to perform, especially in olfaction, for several reasons. First, ORs, like many GPCRs, are difficult to express in their non-native contexts and often require specialized accessory factors and signaling proteins to function heterologously^16^. Second, transient transfection must be performed for tens to hundreds of individual cell lines each time an assay is performed. Thus, experimental protocols for such multiplexed screens are expensive, labor intensive, and often carried out in a low-throughput manner. Using stable lines would alleviate these burdens, but building stable OR reporter lines is challenging and has only worked in two reported cases^17,18^.

Here we report a new high-throughput screen to characterize small molecule libraries against mammalian OR libraries in multiplex. To do this, we developed both a stable cell line capable of functional OR expression (ScL21) and a multiplexed reporter for OR activity (Fig 1a, Supplementary Fig. 1). Activation of each OR leads to the expression of a reporter transcript with a unique 15-nucleotide barcode sequence. Each barcode identifies the OR expressed in that cell; this enables OR activation to be measured by quantifying differential barcode expression with RNA-seq. This technology enables the simultaneous profiling of a single chemical’s activity against a library of receptors in a single well.

**Figure 1.**
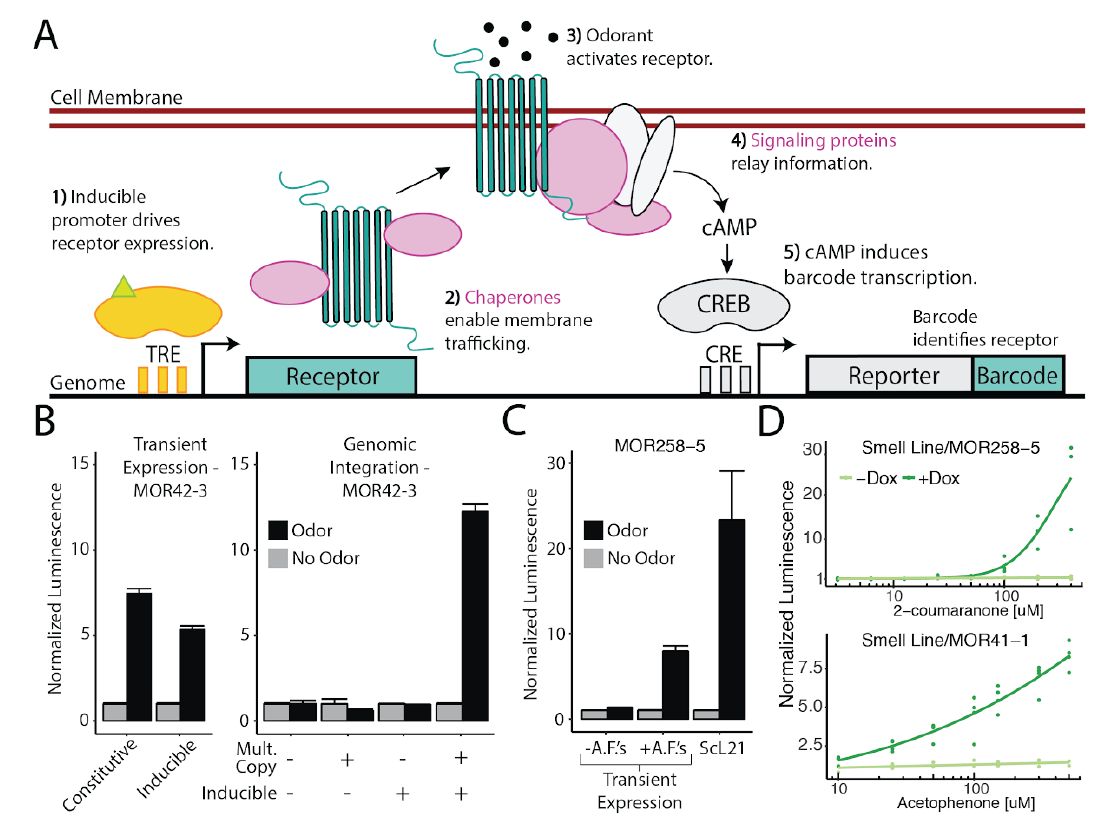
A Genomically Integrated Synthetic Circuit Allows Screening for Mammalian Olfactory Receptor Activation. **A.** Schematic of the synthetic circuit for stable OR expression and function in an engineered HEK293T cell line (ScL21). Heterologous accessory factors expressed include (pink): RTP1S, RTP2, G_αolf_, and Ric8b. **B.** MOR42-3 reporter activation expressing the receptor transiently (left) or genomically integrated (right) at varying copy number, under constitutive or inducible expression in HEK293T cells. **C.** MOR258-5 reporter activation with/without accessory factors (A.F.s), RTP1S and RTP2, transiently coexpressed in HEK293T cells compared to stable receptor expression in ScL21. **D.** Reporter activation response curves for MOR258-5 and MOR41-1 genomically integrated in ScL21.

We first engineered a stable cell line, ScL21, capable of functionally expressing ORs and responding to odorant stimuli by transcribing an RNA barcode. First, we found that multi-copy integration and inducible receptor expression are both essential for reporter activation, but individually neither of these features is sufficient to generate a response (Fig. 1b, Supplementary Fig. 2a, b, c). Then, to allow larger OR repertoires to be assayed, we added features known to improve OR function^16,19,20^. We stably integrated a pool of 4 accessory factors at multi-copy under inducible expression: G_αolf_ and Ric8b for signal transduction, and RTP1S and RTP2 to promote surface expression (Fig. 1c, Supplementary Fig. 2d, Supplementary Fig. 1). To select a single line for further use, we isolated clones and screened for robust activation of two ORs known to require accessory factors to function heterologously (Supplementary Fig. 2e). In addition, we then incorporated protein trafficking tags previously shown to increase surface expression^21,22^, included DNA insulator sequences to reduce background reporter activation, optimized the cAMP response element (CRE) to improve reporter signal, and combined these improvements into a single transposable vector to speed cell line development (Supplementary Fig. 3). We validated our system on two murine ORs with known ligands, and observed induction- and dose-dependent activation (Fig. 1d).

To pilot the platform, we chose 42 murine ORs with both known and unknown chemical specificities and created a library of OR-expressing cell lines. To create the individual cell lines, we first cloned and mapped the ORs to their corresponding barcodes and transposed the plasmids individually into the genomes of HEK293T cells^23^. After selection we pooled the cell lines together, generating assay-ready libraries for repeated testing (Fig. 2a). Unlike a luciferase reporter assay, each well contains the entire OR library and a single chemical’s activity is measured against the entire library of ORs in a single well. We plated the cell library in 6-well culture dishes and screened odorants known to activate ORs in our library (Supplementary Fig. 4a); all but 3 ORs passed quality filtering to obtain reliable estimates of activation (See Methods). Analysis of the sequencing readout recapitulated previously identified odorant-receptor pairs^12^, and chemical mixtures appropriately activated multiple ORs (Supplementary Fig. 4a). We found the assay was robust to nuisance chemicals such as the adenylate cyclase stimulator, forskolin, which non-specifically induces barcode transcription independent of the OR each cell expresses. This is likely because our library-based approach measures the relative activation of ORs to each other, normalizing any global effects due to off-target reporter activation.

**Figure 2.**
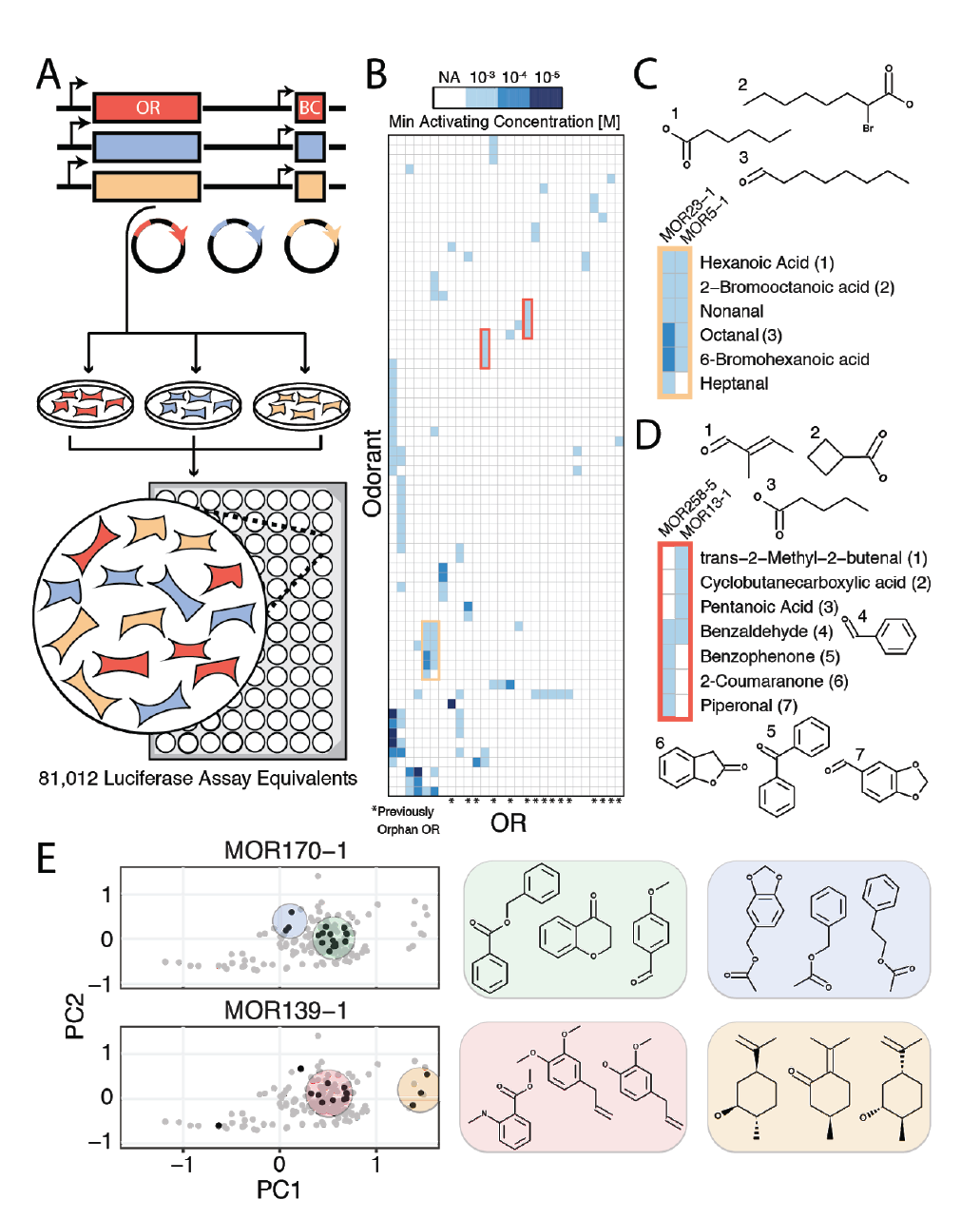
Large-Scale, Multiplexed Screening of Olfactory Receptor-Odorant Interactions. **A.** Experimental workflow for OR library generation and high-throughput screening. To perform assay, we cloned OR genes and barcodes into plasmids, engineered cell lines via individual transposition of plasmids, pooled cell lines and performed screen in 96 well plates. We assayed the equivalent of 81,012 wells of a screen where interactions are tested individually. **B.** Heatmap of interactions from the screen clustered by odorant and receptor responses, and shaded by the minimum activating odorant concentration that triggered reporter activity. Only ORs and chemicals that registered at least one interaction are shown. **C.** Chemical names and structures for odorants that activate MOR23-1 and MOR5-1. **D.** Chemical names and structures for odorants that activate MOR258-5 and MOR13-1. **E.** Chemical hits identified for MOR170-1 and MOR139-1 (black) mapped onto a PCA projection of the chemical space of our odorant panel (grey). Shaded areas highlight hits that cluster together in chemical space.

Next, we adapted the platform for high-throughput screening in 96-well format. To decrease reagent cost and assay time, we optimized an in-lysate reverse transcription protocol and used dual indexing to uniquely link barcode reads to the correct well once samples were mixed for sequencing (see Methods). With these improvements, the assay is able to recapitulate dose-response curves for known odorant-receptor pairs (Supplementary Fig. 4b). We observed reproducible results between identically treated but biologically independent wells (Supplementary Fig. 5 and 6).

We subsequently screened 181 odorants with both known and unknown receptor specificity at three concentrations in triplicate against the 39-member OR cell library, or 81,012 wells if each combination had been tested individually including controls (Fig. 2a and Supplementary Tables 1 and 2). Each 96-well plate in the assay contained independent positive control odorants and solvent (DMSO) for normalization (Supplementary Fig. 6). We used a generalized linear model to determine differentially responsive ORs (see Methods)^24^. We found 112 positive OR-odorant interactions (out of >7,000 combinations), of which 79 are novel, and 24 that target 15 orphan receptors (Benjamini-Hochberg corrected FDR = 1%; Fig. 2b, Supplementary Fig. 7a, and Supplementary Table 3)^25^. Overall, 28 of 39 receptors were activated by at least one odorant, and 67 of 181 odorants activated at least one OR (Supplementary Table 3).

We compared results to a previous study and analyzed individual interactions in a different context. First, we chose 36 interactions with at least 1.2-fold induction to retest individually in a previously developed transient OR activation system^26^ (Supplementary Fig. 8). Of the 27 interactions called as hits at an FDR of 1 %, 20 of them replicated in this orthogonal system (Supplementary Fig. 7b). Notably, some of the seven interactions which did not replicate in this orthogonal system, look like true hits. For instance, our assay registered two hits for MOR19-1 with high chemical similarity (methyl salicylate and benzyl salicylate), suggesting they are likely not false positives (Supplementary Fig. 8). Additionally, three of nine interactions not passing the 1% FDR threshold showed activation in the orthogonal assay, indicating that a conservative FDR threshold likely generated some false negatives. A previous large-scale OR deorphanization study used some of the same receptors and chemicals, and we found that 9/12 of their reported interactions with EC_50_ below 100μΜ were also detected in our platform, though we did not identify most of the previous low affinity interactions^12^ (Supplementary Fig. 9). Conversely, we also detected 14 positive interactions absent from the previous study. Finally, our assay replicated the vast majority of non-interacting odorant-OR pairs (493/507).

Using the data generated by this high throughput assay, we found that chemicals with similar features activate the same ORs, including those receptors we deorphanize in this study (Fig. 2c). For example, the previously orphan MOR19-1 has clear affinity for the salicylate functional group, while MOR13-1 is activated by four chemicals with hydrogen bond accepting groups attached, and in three cases, to stiff non-rotatable scaffolds (Supplementary File 1). We also detect ORs with partial overlap in chemical specificity; MOR13-1 detects compounds with terminal carbonyls while MOR258-5 detects cyclic conjugated molecules (Fig. 2d). Benzaldehyde, an intermediate size carbonyl, activates both ORs.

To more systematically understand how chemical similarity relates to receptor activation, we used a recently developed molecular autoencoder^27^ to computationally map each tested chemical onto a ~292-dimensional continuous latent space with nearly lossless compression (Supplementary Fig. 10). In this space, hits for 11/17 multi-hit receptors cluster together across the first two principal components (Supplementary Fig. 11). For instance, MOR5-1 ligands cluster together and 10 out of 13 aliphatic aldehydes or carboxylic acids with >5 carbons activate the receptor (Fig. 2c, Supplementary File 1). Interestingly, projection onto the chemical latent space also highlights the instances where ORs are sensitive to several distinct sets of chemicals (Fig. 2e). For example, MOR139-1 is activated by compounds that belong to two distinct latent space clusters: one with benzene rings and the other with cyclohexane rings, hinting at the selective features of these odorants. Similarly, MOR170-1 exhibits a broad activation pattern: this receptor responds to ~50% of all odorants in our panel that contains both a benzene ring, and either a carbonyl or ether group. Most of these hits form a cluster in the latent space, with the exception of the acetate compounds in a separate cluster. Understanding the global chemical space that activates each OR establishes the groundwork for the prediction of novel odorant-OR interactions.

We anticipate that this platform can be scaled to test the 396-member human OR repertoire and comprehensively define OR response to any odorant of interest. The approximate cost per well is on par with existing assays, but per receptor-ligand interaction interrogated, multiplexing dramatically reduces cost and labor. Our incomplete understanding for how ligands^28^, drugs^1^, hormones, natural products^29^ and odors^12^ interact with potential cellular targets limits our ability to rationally develop new molecules to modulate receptor activity. Multiplex methods like this platform offer a scalable solution to generate large-scale datasets that will help guide both empirical and algorithmic efforts to better dissect the complex interactions between small molecules and biological targets^30^.

## Acknowledgements

We thank the Kosuri Lab and Joshua Bloom for helpful discussions, the UCLA Broad Stem Cell Research Center Sequencing Core, and the Technology Center for Genomics and Bioinformatics for providing next-generation sequencing. We thank Mengjue Ni, Jeong Hoon Ko, and Aaron Cooper for expert technical assistance. We thank Jason Kakoyiannis for providing perfumes as a gift. Funding provided by National Science Foundation, Brain Initiative (1556207 to S.Kand 1556207/151801 to H.M.), the Jane Coffin Child Foundation (R.J.), Ruth L. Kirschstein National Research Service Award (GM007185 to N.L.), the USPHS National Research Service Award (5T32GM008496 to E.J.), the NIH (DC014423 to H.M.) and UCLA.

E.J., R.J., N.L. and S.K. consult for and hold equity in Octant Inc. to which patents based on this work have been licensed. All other authors declare that they have no competing interests.

## References

1. Roth B. L., Sheffler D. J. & Kroeze W. K. Magic shotguns versus magic bullets: selectively non-selective drugs for mood disorders and schizophrenia. Nat. Rev. Drug Discov. 3, 353–359 (2004).

2. Reddy A. S. & Zhang S. Polypharmacology: drug discovery for the future. Expert Rev. Clin. Pharmacol. 6, 41–47 (2013).

3. Fang J., Liu C., Wang Q., Lin P. & Cheng F. In silico polypharmacology of natural products. Brief. Bioinform. (2017). doi:10.1093/bib/bbx045

4. Anighoro Α., Bajorath J. & Rastelli G. Polypharmacology: challenges and opportunities in drug discovery. J. Med. Chem. 57, 7874–7887 (2014).

5. Malnic B., Hirono J., Sato T. & Buck L. B. Combinatorial receptor codes for odors. Cell96, 713–723 (1999).

6. Buck L. & Axel R. A novel multigene family may encode odorant receptors: a molecular basis for odor recognition. Cell 65, 175–187 (1991).

7. Hauser A. S., Attwood M. M., Rask-Andersen M., Schiöth H. B. & Gloriam D. E. Trends in GPCR drug discovery: new agents, targets and indications. Nat. Rev. Drug Discov. 16, 829–842 (2017).

8. Niimura Y., Matsui A. & Touhara K. Extreme expansion of the olfactory receptor gene repertoire in African elephants and evolutionary dynamics of orthologous gene groups in 13 placental mammals. Genome Res. 24, 1485–1496 (2014).

9. Peterlin Z., Firestein S. & Rogers M. E. The state of the art of odorant receptor deorphanization: a report from the orphanage. J. Gen. Physiol. 143, 527–542 (2014).

10. Lu M., Echeverri F. & Moyer Â. D. Endoplasmic reticulum retention, degradation, and aggregation of olfactory G-protein coupled receptors. Traffic 4, 416–433 (2003).

11. Bushdid C, de March Ñ Α., Matsunami H. & Golebiowski J. Numerical Models and In Vitro Assays to Study Odorant Receptors, in Olfactory Receptors: Methods and Protocols (eds. Simoes de Souza F. M. & Antunes G.) 77–93 (Springer New York, 2018).

12. Saito H., Chi Q., Zhuang H., Matsunami H. & Mainland J. D. Odor coding by a Mammalian receptor repertoire. Sci. Signal. 2, ra9 (2009).

13. Mainland J. D. et al. The missense of smell: functional variability in the human odorant receptor repertoire. Nat. Neurosci. 17, 114–120 (2014).

14. Botvinik A. & Rossner M. J. Linking cellular signalling to gene expression using EXT-encoded reporter libraries. Methods Mol. Biol. 786, 151–166 (2012).

15. Galinski S., Wiehert S. P., Rossner M. J. & Wehr M. C. Multiplexed profiling of GPCR activities by combining split TEV assays and EXT-based barcoded readouts. Sci. Rep. 8, 8137 (2018).

16. Zhuang H. & Matsunami H. Synergism of accessory factors in functional expression of mammalian odorant receptors. J. Biol. Chem. 282, 15284–15293 (2007).

17. Cook B. L, Ernberg Ê. E., Chung H. & Zhang S. Study of a synthetic human olfactory receptor 17-4: expression and purification from an inducible mammalian cell line. PLoS One 3, e2920 (2008).

18. Belloir C, Miller-Leseigneur M.-L, Neiers F., Briand L. & Le Bon A.-M. Biophysical and functional characterization of the human olfactory receptor OR1A1 expressed in a mammalian inducible cell line. Protein Expr. Purif. 129, 31–43 (2017).

19. Saito H., Kubota M., Roberts R. W., Chi Q. & Matsunami H. RTP family members induce functional expression of mammalian odorant receptors. Cell 119, 679–691 (2004).

20. Von Dannecker L. E. C., Mercadante A. F. & Malnic B. Ric-8B, an olfactory putative GTP exchange factor, amplifies signal transduction through the olfactory-specific G-protein Galphaolf. J. Neurosci. 25, 3793–3800 (2005).

21. Shepard B. D., Natarajan N., Protzko R. J., Acres O. W. & Pluznick J. L. A cleavable N-terminal signal peptide promotes widespread olfactory receptor surface expression in HEK293T cells. PLoS One 8, e68758 (2013).

22. Krautwurst D., Yau Ê. W. & Reed R. R. Identification of ligands for olfactory receptors by functional expression of a receptor library. Cell 95, 917–926 (1998).

23. Li X. et al. piggyBac transposase tools for genome engineering. Proc. Natl. Acad. Sci. U. S. A. 110, E2279–87 (2013).

24. McCarthy D. J., Chen Y. & Smyth G. K. Differential expression analysis of multifactor RNA-Seq experiments with respect to biological variation. Nucleic Acids Res. 40, 4288–4297 (2012).

25. Benjamini Y. & Hochberg Y. Controlling the False Discovery Rate: A Practical and Powerful Approach to Multiple Testing. J. R. Stat. Soc. Series Â Stat. Methodol. 57, 289–300 (1995).

26. Zhuang H. & Matsunami H. Evaluating cell-surface expression and measuring activation of mammalian odorant receptors in heterologous cells. Nat. Protoc. 3,1402–1413 (2008).

27. Gómez-Bombarelli R. et al. Automatic Chemical Design Using a Data-Driven Continuous Representation of Molecules. ACS Cent Sci 4, 268–276 (2018).

28. Antebi Y. E. et al. Combinatorial Signal Perception in the BMP Pathway. Cell 170,1184–1196.e24 (2017).

29. Harvey A. L, Edrada-Ebel R. & Quinn R. J. The re-emergence of natural products for drug discovery in the genomics era. Nat. Rev. Drug Discov. 14, 111–129 (2015).

30. Colwell L. J. Statistical and machine learning approaches to predicting protein-ligand interactions. Curr. Opin. Struct. Biol. 49, 123–128 (2018).

